# A novel dual antibody staining assay to measure estrogen receptor transcriptional activity

**DOI:** 10.1101/2020.04.14.021782

**Authors:** Freek van Hemert, Christa Dam-de Veen, Sil Konings, John van der Ven, Anja van de Stolpe

**Affiliations:** Precision Diagnostics, Philips Research, Eindhoven, The Netherlands

## Abstract

Activity of the canonical estrogen receptor (ER) pathway is equivalent to functional activity of the nuclear ER transcription factor. To assess transcriptional activity of ER, for biomedical research and diagnostic purposes ER monoclonal antibodies are routinely used to identify nuclear ER staining in cells and tissue samples, however it remained unclear whether this is sufficiently predictive for transcriptional activity of ER, and thus for ER pathway activity. Using ER positive breast cancer cell lines (MCF7 and T47D) in which the transcriptional activity status of ER was quantified using an mRNA based ER pathway activity assay, the relation between ER activity and nuclear ER staining with ER monoclonal antibodies (MoAb) was investigated. While the presence of ER in the cell nucleus is a prerequisite for ER activity, it was not predictive for ER transcriptional activity, confirming earlier findings. There were remarkable differences in behaviour of the used MoAbs: EP1 and 1D5 MoAbs showed reduced nuclear staining when ER was transcriptionally active, while staining with H4624 MoAb was independent of ER activity. To improve discrimination between active and inactive nuclear ER based on ER staining, a method was developed which consists of dual ER MoAb immunofluorescent staining, followed by generation of a digital image with a standard digital pathology scanner, and application of a cell nucleus detection algorithm and *per cell* calculation of the nuclear H4624/EP1 fluorescence intensity ratio, where a high H4624/EP1 ratio predicts an active ER. In this method the EP1 MoAb can in principle be replaced by the 1D5 MoAb. We hypothesize that the EP1 and 1D5 monoclonal antibodies (MoAb) recognize an ER epitope which becomes hidden upon transcriptional activation of ER, while the H4624 MoAb binds an ER epitope which remains accessible when ER is activated. The method is expected to be of value to better assess ER activity by means of staining.

## Introduction

The estrogen receptor (ER) signal transduction pathway plays an important role in physiological processes, for example in the menstrual cycle, and in the pathophysiology of a variety of diseases, such as breast cancer. Simplified, the signalling pathway consists of an estrogen ligand which specifically binds to the estrogen receptor, resulting in receptor dimerization and recruitment of co-activator molecules, and initiation of transcription of target genes of ER, that is, genes containing an estrogen response element in their promoter region. The produced target gene mRNAs are translated into proteins which change the behaviour of the cell as response to the estrogen signal, e.g. cell division or differentiation. Other signalling pathways may crosstalk with the ER pathway, increasing the complexity of this cellular mechanism. Being able to measure the activity of the ER signalling pathway is important, not only to unravel complex ER signalling and intracellular crosstalk mechanisms, but also for drug development, and in diagnostics, for example to subtype patients with breast cancer and decide on hormonal therapy.

ER antibody staining is frequently used for this purpose and positive nuclear ER staining is considered indicative of an active ER pathway. However it is unclear how nuclear ER staining relates to ER transcriptional activity. As an illustration of the problem, in general up to half of breast cancer patients selected for hormonal therapy based on positive ER staining do not respond, suggesting that ER expression does not necessarily imply an active ER pathway in cancer tissue ^1^,^2^.

To address this issue, we have previously developed an assay which allows quantitative measurement of ER pathway activity based on measuring ER target gene mRNA levels followed by inferring ER pathway activity using a knowledge-based Bayesian computational model ^1^. Evaluation of this ER pathway assay on a large number of ER positive and negative breast cancer samples and more recently on three independent clinical patient cohorts treated with neoadjuvant hormonal therapy, provided evidence that nuclear ER staining is required, but not sufficient, for ER pathway activity ^1^,^3^.

Since it remains important to be able to assess ER pathway activity on cell and tissue samples without the need to disrupt the cells to extract RNA, we here describe development of a fluorescent dual antibody ER staining assay, which improves prediction of the ER transcriptional activity state in cells on a cytology or tissue slide. The staining assay is expected to provide complementary information to the mRNA-based ER pathway activity assay, for example spatial information on cellular ER pathway activity within an intact tissue architecture.

## Methods

### Cell culture

Human epithelial cell breast cancer lines, MCF7 (HTB-22, ER+), T47D (HTB-133, ER+), and MDA-MB-231 (HTB-26, ER-), were purchased from ATCC (Washington, USA). Cells were cultured at 37°C, 5% CO_2_, 95% humidity, in appropriate culture medium (Gibco, Carlsbad, California, USA). MCF7 and MDA-MB-231 were cultured in Dulbecco’s Modified Eagles Medium (DMEM) and T47D was cultured in Roswell Park Memorial Institute medium (RPMI). Medium was supplemented with 1% penicillin/streptomycin, 10% fetal bovine serum (FBS) and 1% Glutamax (Gibco, Carlsbad, California, USA). To generate ER pathway-inactive and ER pathway-active breast cancer cells, the cells were first deprived from estrogen for two days and subsequently stimulated with estradiol, according to Katzenellenbogen et al. and as described before^4^,^1^. In brief, prior to estradiol stimulation, MCF7 cells were cultured for 48 hours in phenol red-free culture medium, supplemented with 10% heat-inactivated and charcoal-treated fetal bovine serum (FBS) and 1% Glutamax. Subsequently, cells were stimulated with estradiol (E2, 10 nM, Sigma-Aldrich, Saint Louis, USA) or vehicle (DMSO, Sigma-Aldrich, Saint Louis, USA) for 16 hours. These experiments were performed by BioDetection Systems (Amsterdam, The Netherlands). For fluorescent ER protein staining experiments the same estrogen deprivation and stimulation protocol was followed, only cells were cultured on fibronectin-coated coverslips for various timepoints as indicated. For the dual fluorescent staining assays, staining was performed at 0 and 30 minutes.

### Immunofluorescent ER staining

Cells were fixated with 4% pH neutral buffered formaldehyde (Boom BV, Meppel, The Netherlands) for 15 minutes, permeabilized and blocked with 0.2% Triton X-100/1% bovine serum albumin (BSA)/PBS solution for 20 minutes at room temperature (RT). Coverslips were rinsed, incubated with primary antibody for one hour in a humidified environment at RT, rinsed, incubated with secondary antibody for 30 minutes in a humidified and dark environment at RT, rinsed, and finally mounted on microscope slides with Vectashield Hard Set mounting medium containing DAPI (H-1500, Vector laboratories, Burlingame, California, USA).

Primary antibodies used were monoclonal mouse anti-human estrogen receptor-α 1 (1D5), monoclonal mouse anti-human estrogen receptor-α 2 (EP1), monoclonal rabbit anti-human estrogen receptor-α (H4624) and polyclonal rabbit anti-human estrogen receptor-α (H-184). EP1 and 1D5 MoAbs are used in routine clinical pathology. Secondary antibodies were goat anti-mouse ATTO-555 (1/200 dilution, Invitrogen, Carlsbad, California, USA) and/or goat anti-rabbit ATTO-633 (1/200 dilution, Invitrogen Carlsbad, California, USA) (for details, see Supplemental Table 1). For dual antibody ER staining, primary monoclonal antibodies (MoAb) from mouse and rabbit were used in combination. The ER negative MDA-MB-231 cell line was used as negative control for staining with ER MoAbs.

### Measurement of functional ER pathway activity

Following deprivation and stimulation with estradiol, RNA was isolated using the Nucleospin RNA isolation kit (Macherey-Nagel)(BioDetection Systems, Amsterdam). Affymetrix HGU133Plus2.0 microarray analysis was performed by Service XS (Leiden, The Netherlands). From the Affymetrix microarray expression data the ER pathway activity score was calculated using the ER pathway activity assay, as described before^1^,^3^. In brief, the ER pathway assay infers transcriptional activity of ER from mRNA levels of a number of high evidence ER target genes, in this case measured by Affymetrix microarray. The calculated ER transcriptional activity score is taken as equivalent of the ER pathway activity score. Activity scores are presented on a log2 odds scale^5^,^6^.

### Cell image analysis and nuclear ER staining quantification algorithm

Stained coverslips were scanned on a 3D Histech Pannoramic MIDI system equipped with a 20x objective and a CIS 3CCD camera. For dual antibody ER staining, fluorescence detection was performed in separate channels. For the DAPI channel 25 millisecond (ms) exposure without digital gain was used, for the TRITC channel 150 ms exposure without digital gain, and for the Cy5 channel 175 ms exposure with digital gain setting 3. Data were stored in MIRAX file format and converted to RGB TIFF files. Algorithms for analysis of digital images and quantification of immunofluorescent staining intensities were developed using MATLAB 2012b (The MathWorks Inc., Natick, MA, 2000, United States). Digital image analysis consisted of: (1) image pre-processing; (2) cell nucleus detection; (3) cell membrane location estimation; and (4) quantification of immunofluorescence staining intensity in detected nuclei. Details are described in Supplementary Methods.

### Fluorescent staining data analysis

For all cells on a sample slide, nuclear immunofluorescence intensity values were calculated and data were analysed and plotted using the Python3 (Python Software Foundation, https://www.python.org/) modules Pandas ^7^, matplotlib ^8^, Seaborn ^9^ and scikit-learn ^10^. Immunofluorescence intensity results were quantitatively compared using intensity histograms.

### Prediction of ER activity from dual ER MoAb immunofluorescent staining data

ER activity status per individual cell was calculated based on comparison between nuclear fluorescent staining intensity of EP1 and H4624 MoAb. H4624 staining intensity represented the total amount of nuclear ER, that is, both functionally active and inactive, while EP1 (or alternatively 1D5) staining intensity represented only the amount of inactive ER. To predict ER activity, H4624 and EP1 fluorescent intensity signals were used to generate a H4624/EP1 staining intensity ratio for each individual cell nucleus. Since the polyclonal H184 ER antibody also stains ER independent of its activity state, a similar ratio to predict ER activity can be calculated by replacing H4624 staining intensity by H184 staining intensity.

This ratio number was used to generate a Receiver Operating Curve (ROC) curve for H4624/EP1 ratio-based prediction of nuclear ER activity. In addition, ROC curves were generated based on the staining intensities of individual antibodies.

## Results

### Nuclear ER staining in ER positive breast cancer cells with an inactive or active ER

We first investigated the relationship between nuclear ER staining and transcriptional ER activity. The ER positive MCF7 and T47D breast cancer cell lines were used as in vitro model system for inactive and active ER pathway. Estrogen deprivation resulted in a low log2 odds ER pathway activity score associated with an inactive ER pathway, while upon stimulation with estradiol the pathway activity score increased, defining this condition as ER pathway active (Fig. 1).

**Figure 1.**
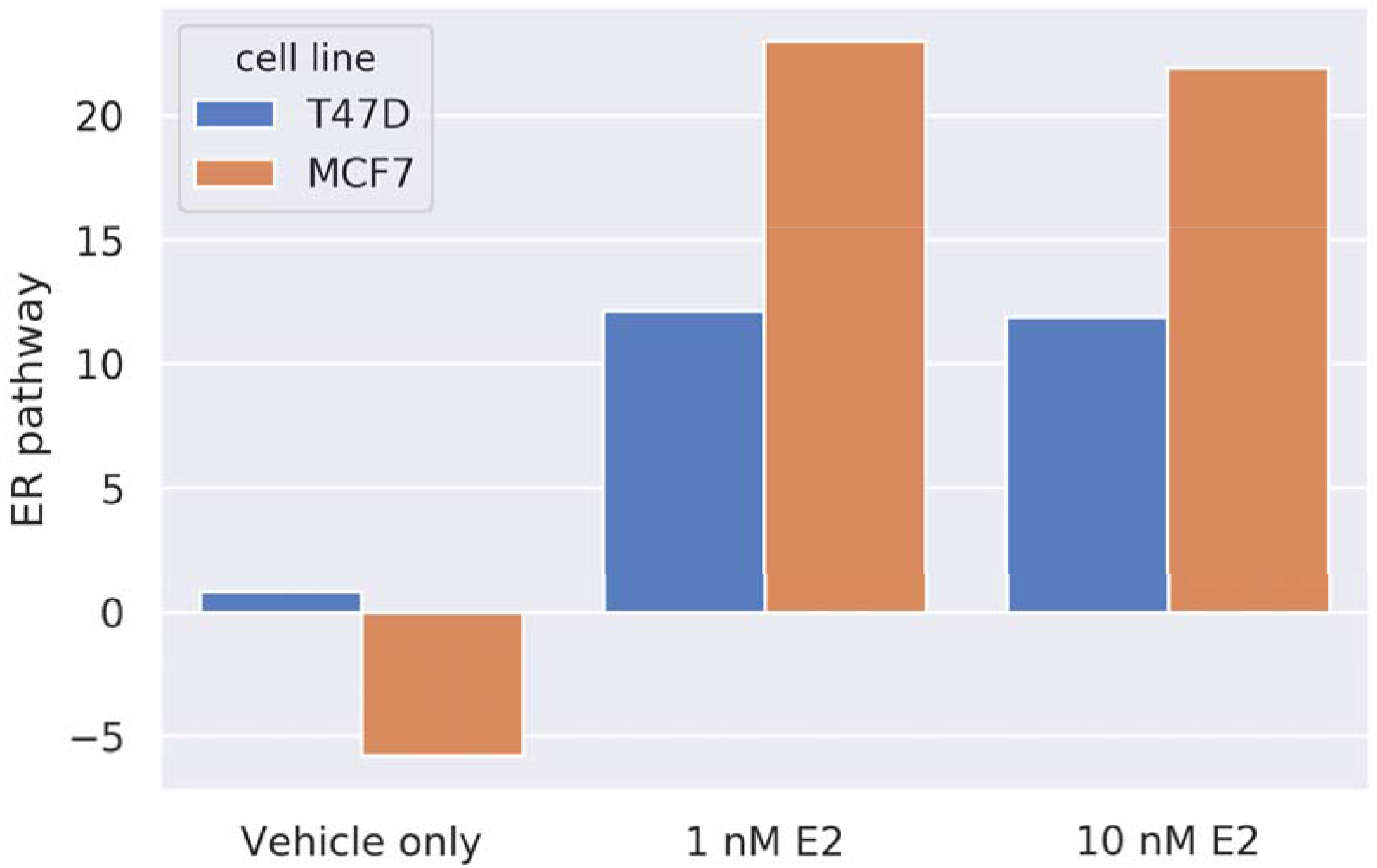
Breast cancer cell lines (T47D, MCF-7) were deprived of estradiol for 48 hours and subsequently stimulated with repectively 1 or 10 nM estradiol for 16 hours. RNA was isolated and ER pathway activity was measured using Affymetrix U133P2.0 microarray expression analysis (see Methods for the used method). ER pathway activity score is indicated as log2 odds of the probability that the ER pathway is active.

Subsequently immunofluorescent ER staining was performed using the EP1 MoAb on MCF7 cells in which the ER pathway was either inactive, or activated by estradiol. EP1 showed specific nuclear ER staining in estrogen-deprived cells with an inactive ER pathway, while nuclear ER staining intensity decreased after activation of the ER pathway with estradiol (Fig. 2). This semi-quantitative analysis suggested that nuclear ER expression was present in the absence of transcriptional activity, while induction of ER transcriptional activity did not increase, but even reduced nuclear ER staining with EP1. Based on this observation we proceeded to quantify the relationship between ER transcriptional activity and immunofluorescent ER staining.

**Figure 2.**
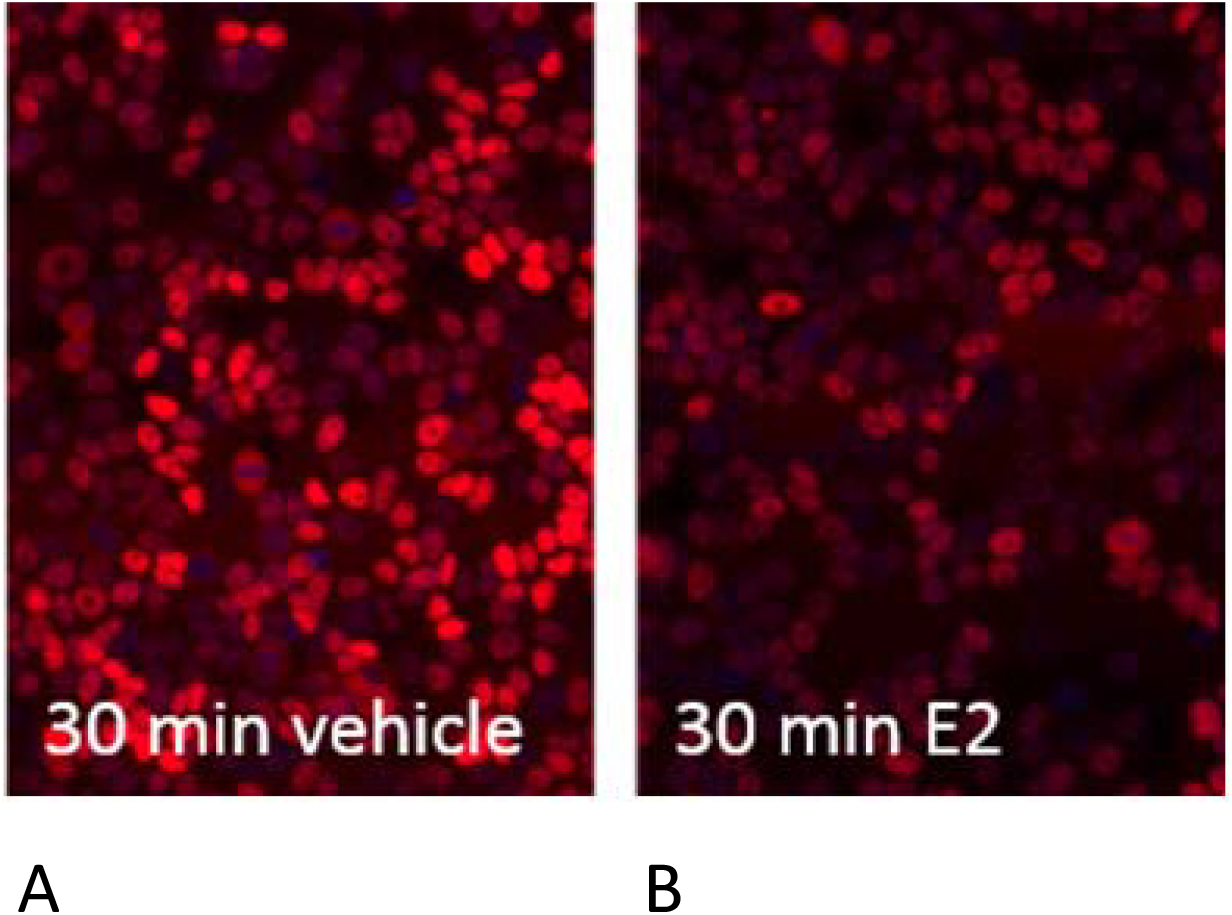
Nuclear ER staining in breast cancer cells with an inactive or active ER pathway. A. E2 depleted MCF7 cells stimulated with vehicle (DMSO) for 30 minutes. B. E2 depleted MCF7 cells stimulated with 10 nM E2 for 30 minutes. Magnification 200x (20x objective).

### Development of an algorithm for quantitative assessment of nuclear ER staining per cell

To enable quantification of the relationship between nuclear ER staining and ER activity, staining with EP1, 1D5, and H4624 MoAbs was performed on MCF7 and T47D cells at different timepoints after estradiol stimulation (5, 10, 30, 60 minutes) (Fig 4A-L). Quantification of immunofluorescent nuclear staining required image scanning and development of nucleus-recognition software. The algorithm for nucleus detection functioned well on scanned slides containing stained cells, as shown in the example slide shown in Fig. 3.

**Figure 3.**
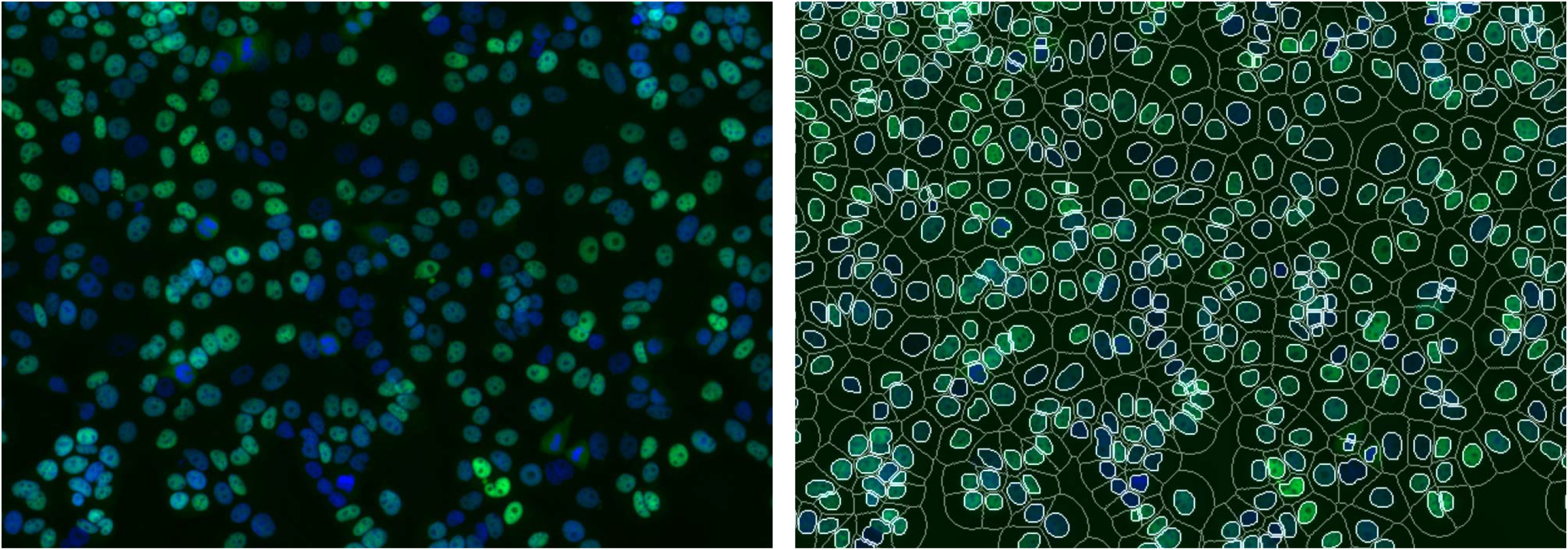
The nucleus detection algorithm reliably detects cell nuclei. A: Cells were cultured on a glass coverslip, routinely fixated and nuclei were stained with DAPI. B: The slide with the coverslip was scanned as described, and nucleus and cytoplasm identified by the nucleus detection algorithm. In blue, DAPI, in Green ER.

**Figure 4.**
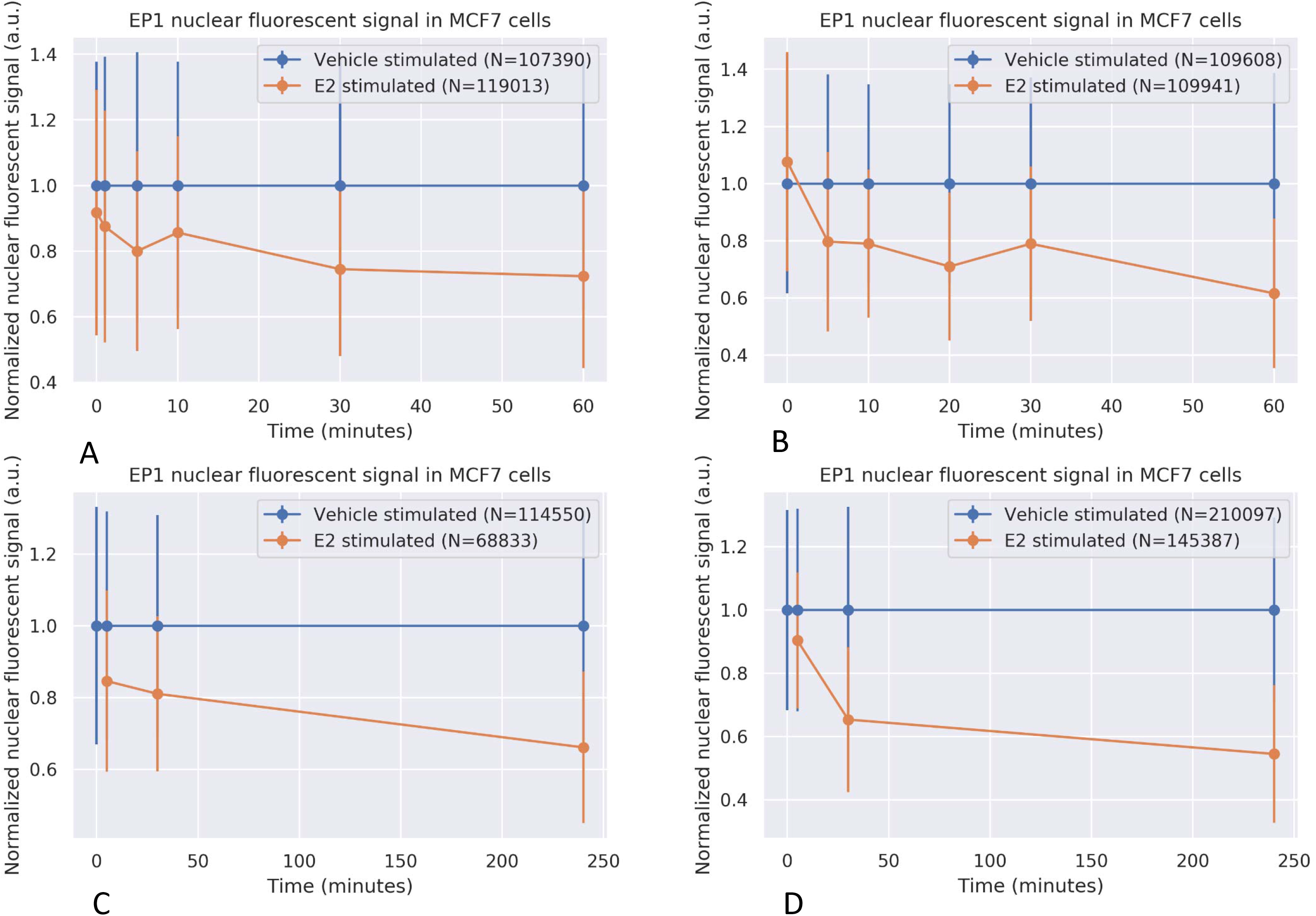
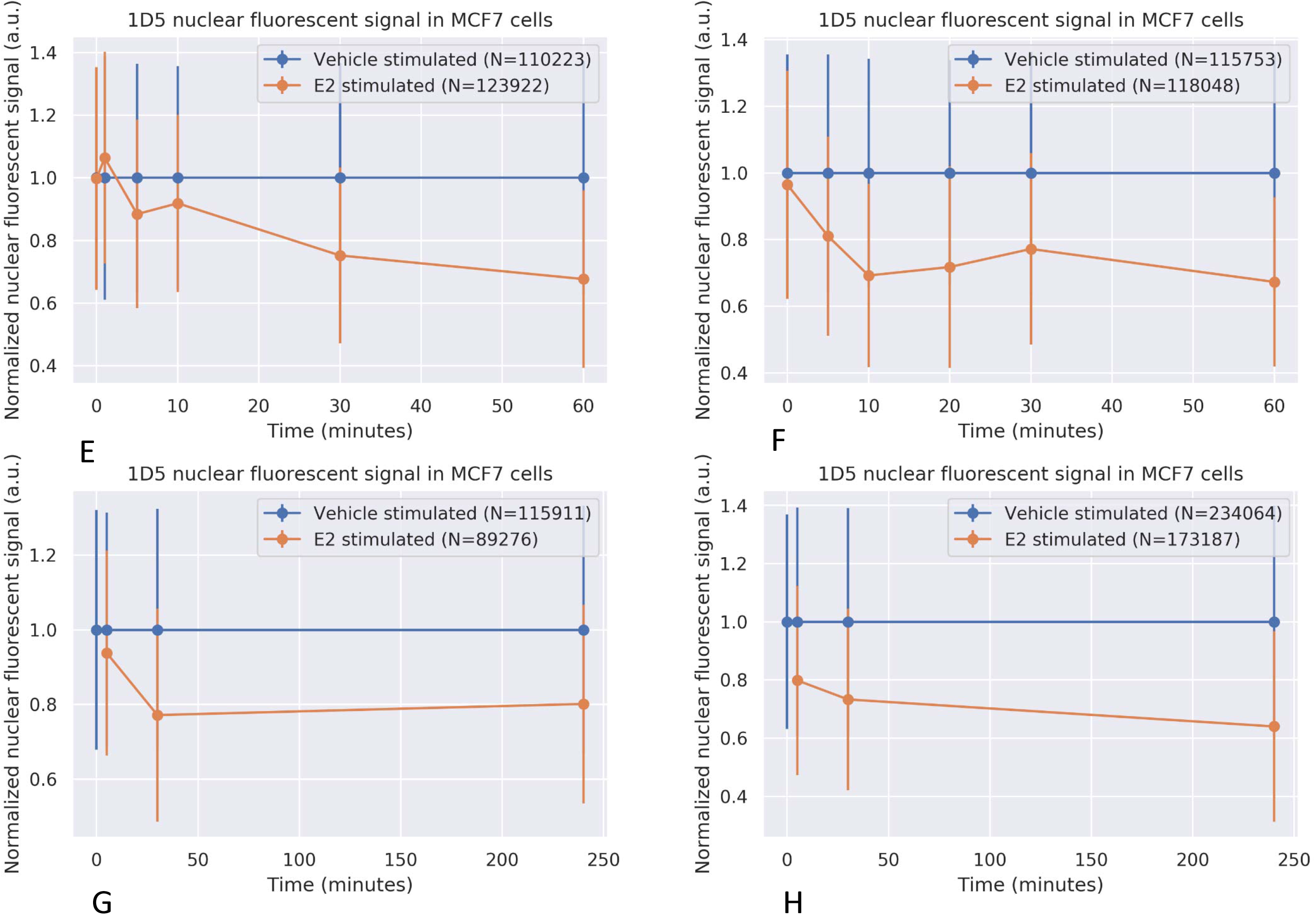
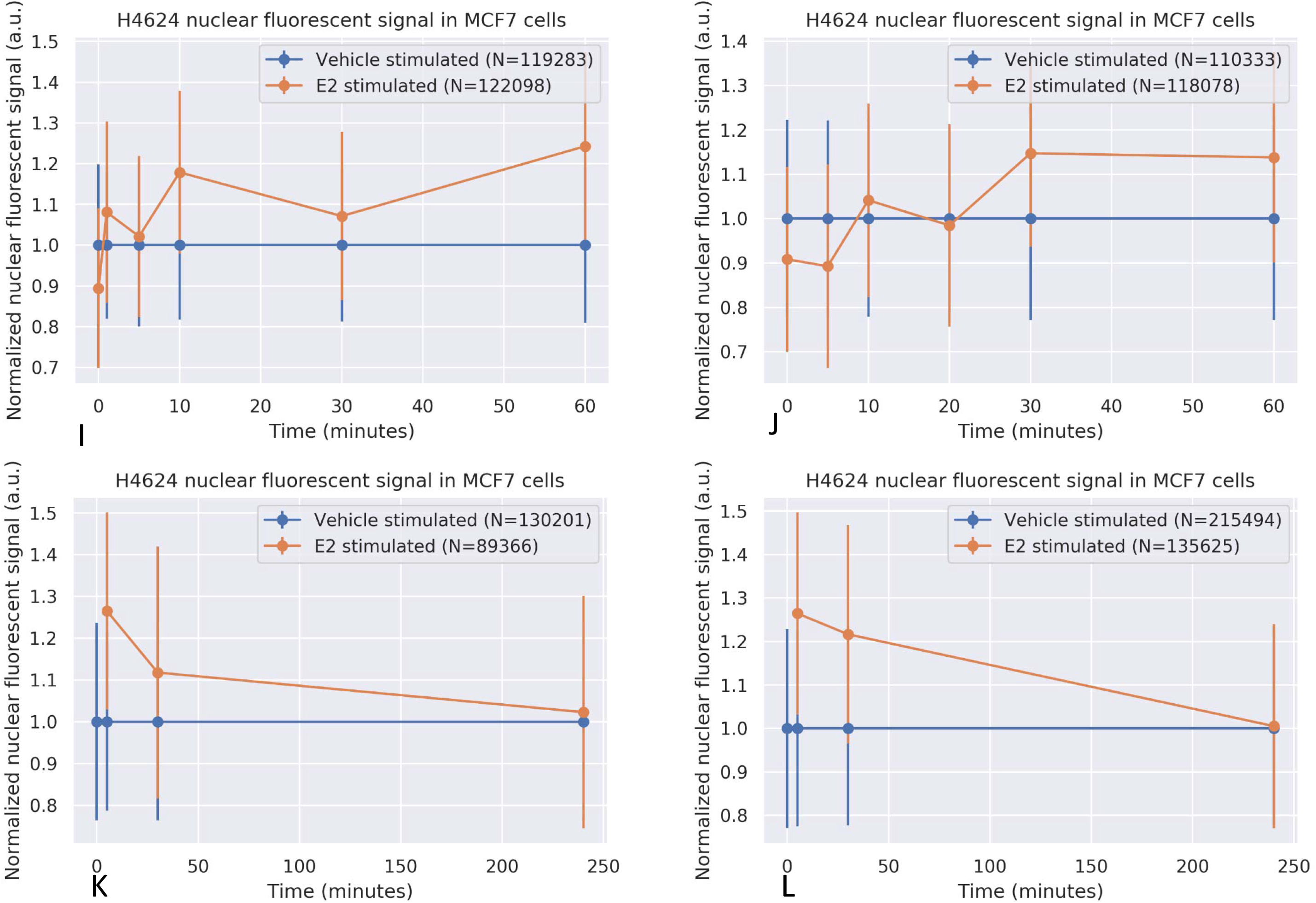
Nuclear ER staining in breast cancer cells with an inactive or active ER pathway. Estrogen-deprived MCF7 cells were either treated with vehicle (blue line) or with 10 nM estradiol (red line). At indicated time points immunofluorescent staining was performed using the indicated MoAb. Nuclear staining intensity was measured and normalized per individual cell, using the nucleus detection+ER-staining algorithm (y-axis). Exposed time is depicted on the X-axis in minutes. For each MoAb 4 independent experiments are depicted. Bars indicate the standard deviation of nuclear staining for a measurement of 10000 to 70000 cells A-D: EP1 MoAb; E-H: 1D5 MoAb; I-L: H4624 MoAb. N indicates the number of analysed cells.

Nucleus detection was used to quantify nuclear fluorescence intensity per cell in a scanned image of a slide and construct histograms from all cells combined. Compared to nuclear ER staining intensity in estrogen-deprived cells, ER staining intensity decreased already within five minutes after estradiol stimulation with either the EP1 or the 1D5 MoAb (Fig. 4A-D and 4E-H).

In contrast nuclear staining intensity with the H4624 MoAb did not decrease after cellular stimulation with estradiol (Fig. 4I-L). On the ER positive cell line T47D similar results were obtained (Supplementary Fig. 1A,B and 2A,B). Use of the polyclonal ER antibody H184 resulted in staining intensity results comparable to those obtained with the H4624 MoAb and independent of ER activity state (Supplementary Fig 2A,B). Cell histograms illustrate in a quantitative manner that EP1 nuclear staining intensity shifted downward upon ER activation, while H4624 staining was not altered (Fig. 5AB). It can also be seen that there was heterogeneity in nuclear ER staining between cells on the analysed slides (Fig. 5AB). The difference between EP1 (or 1D5) and H4624 ER staining signals in a cell appeared to be strongly related to the level of nuclear ER expression: cells with low ER staining, reflecting low ER expression, show no change in staining intensity upon estradiol treatment, while cells with highest ER expression showed the largest reduction in EP1 staining intensity (Fig. 5C).

**Figure 5:**
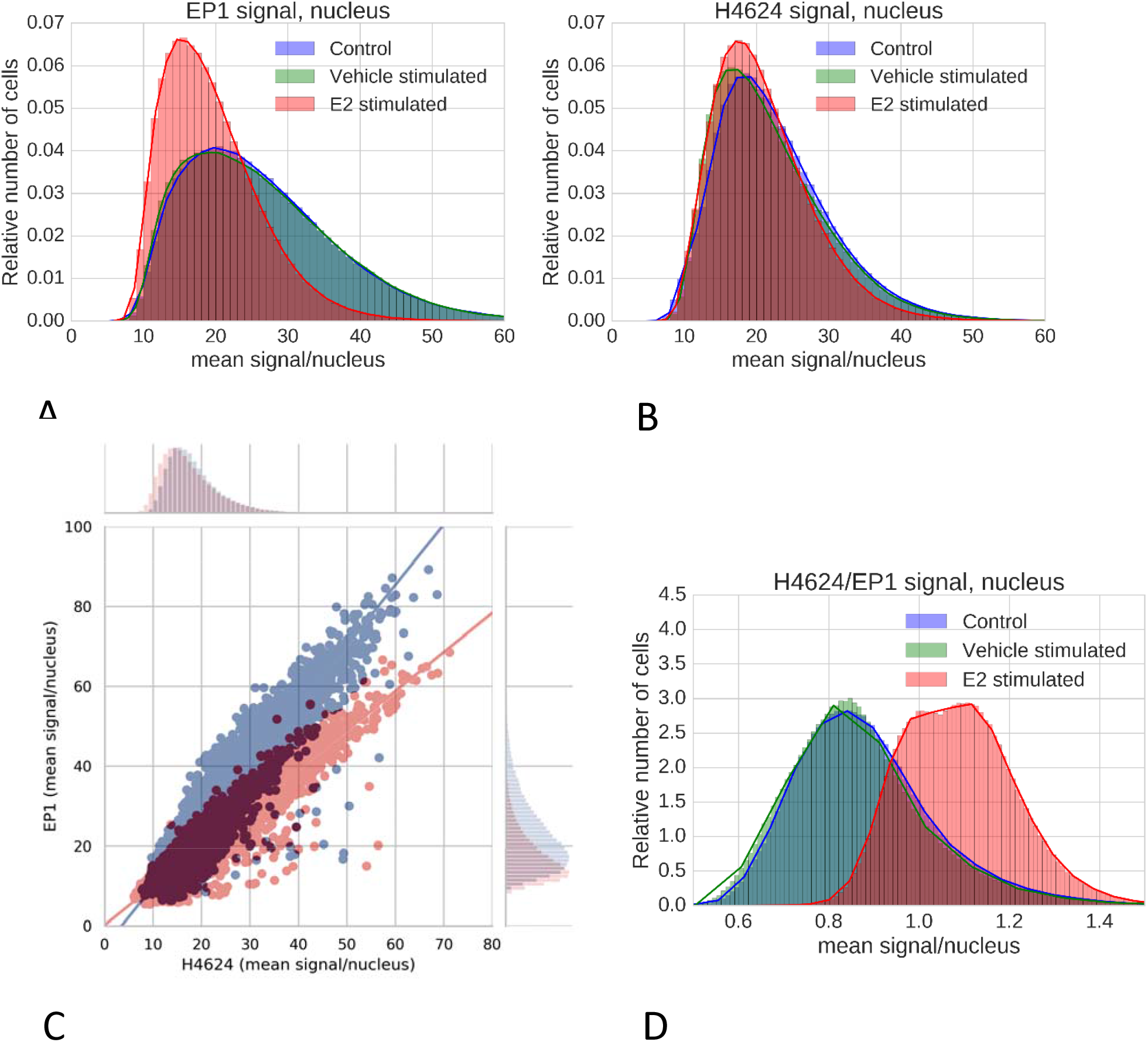
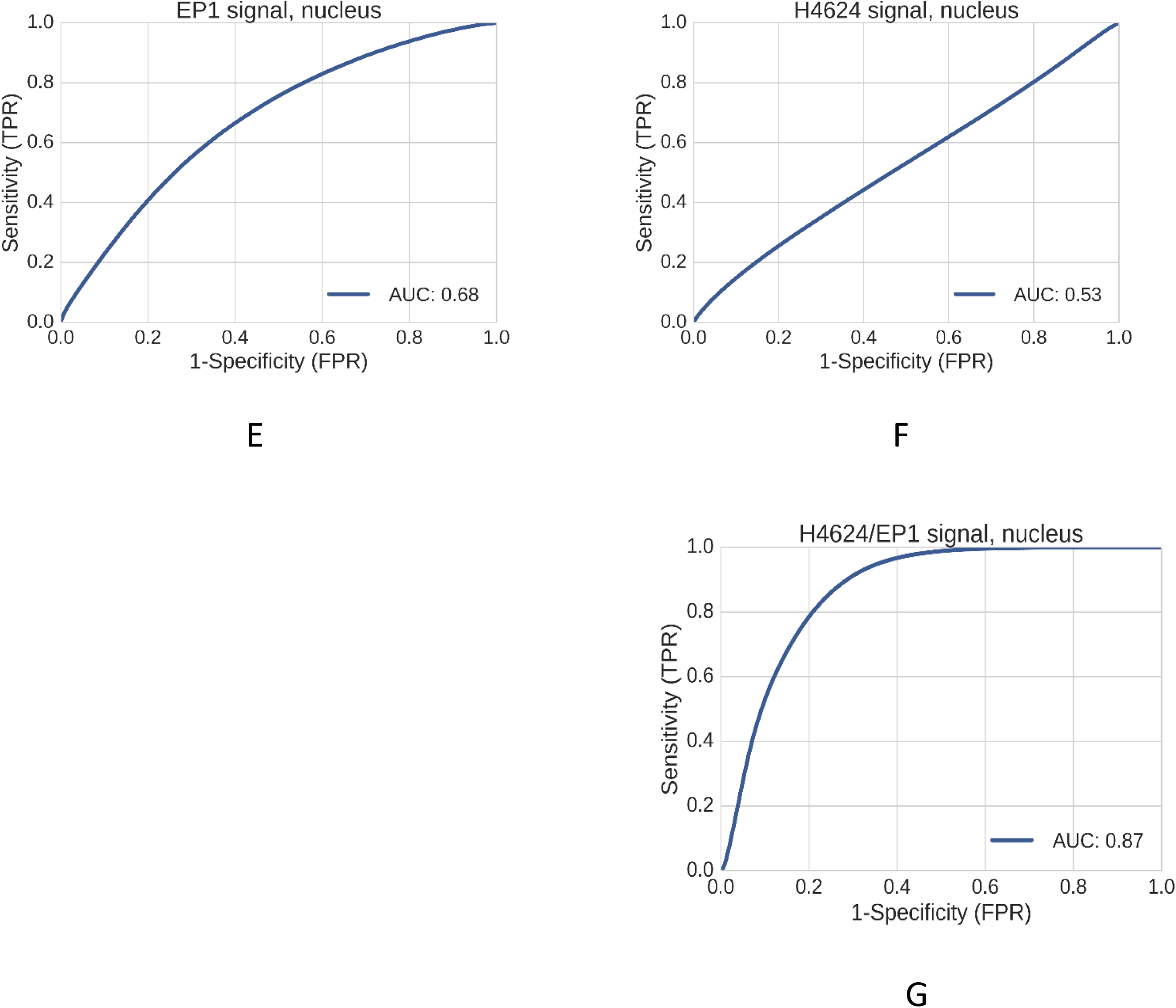
Histograms of mean nuclear fluorescent intensity signal generated using immunofluorescent staining with indicated Moabs in estradiol-deprived MCF7 cells, either treated with vehicle or estradiol for 30 minutes. Blue: E2-depleted MCF7 cells (n = 771444 cells) (control); green: E2 depleted cells treated for 30 minutes with DMSO vehicle (DMSO, n = 758369 cells); red: E2-depleted cells stimulated with 10 nM E2 for 30 minutes (n = 789734 cells). Fluorescence intensity is in Arbitrary Fluorescence Units (AFUs), provided by the digital slide image scanner. N=12 independent experiments, performed in duplo. A. EP1 MoAb; B. H4624 MoAb; C. Correlation between immunofluorescent signal intensity of H4624 and EP1 MoAb. Shifts between the two correlation plots are visualized as corresponding histograms at the top and right sides of the figure quadrant; the shift from *blue* to *red* illustrates per cell the measured decrease in nuclear staining with the EP1 MoAb following cell stimulation with estradiol, relative to nuclear staining obtained with the H4624 MoAb; D.Histograms of the ratio of the mean nuclear fluorescent staining intensity with EP1 and with H4624 MoAb; E. Sensitivity and specificity of the EP1 staining intensity ratio to identify transcriptionally active nuclear ER staining, ROC curve, Area Under the Curve, AUC = 0.68; F. Sensitivity and specificity of the H4624 staining intensity ratio to identify transcriptionally active nuclear ER staining, ROC curve, AUC = 0.53; G. Sensitivity and specificity of the H4624/EP1 staining intensity ratio to identify transcriptionally active nuclear ER staining, ROC curve, AUC = 0.87.

**Figure 6.**
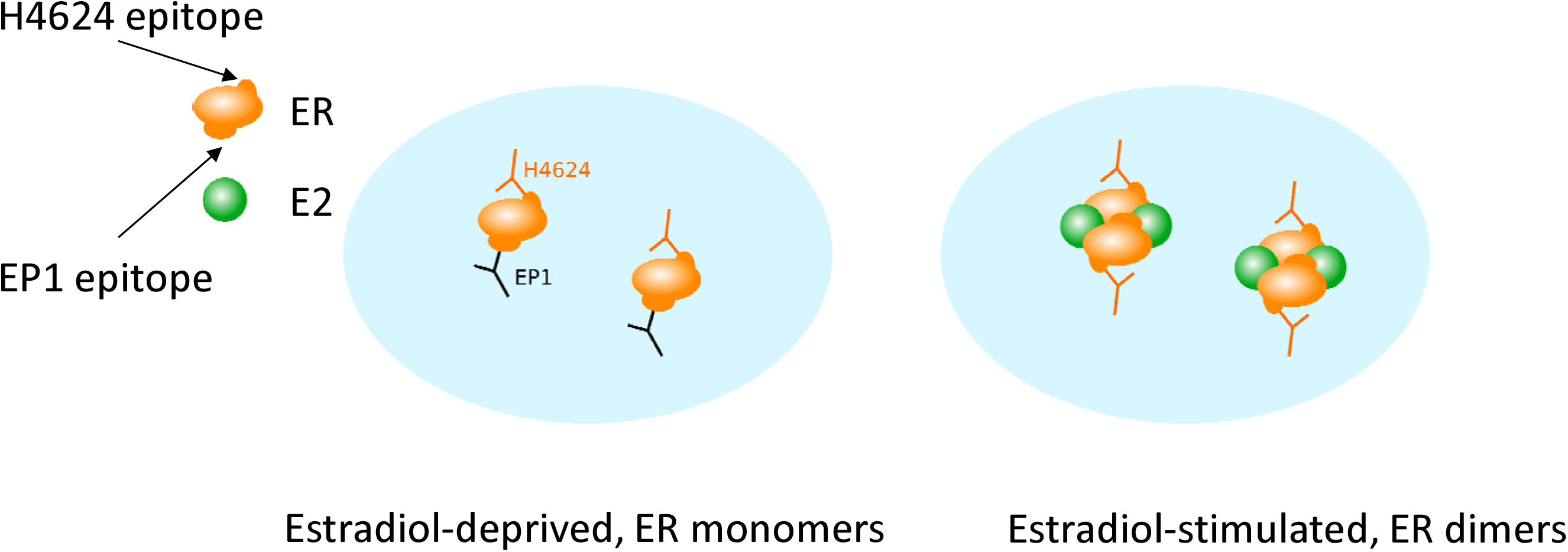
Proposed mechanism of MoAb binding in ER positive MCF7 cells with either a functionally inactive (A) or active (B) ER. A. ER is in its inactive, monomeric state when epitopes for both EP1 and H4624 MoAbs are accessible; B. E2 induces ER dimerization and binding of co-activators, resulting in a transcriptionally active multi-protein ER transcription factor complex. During this process, the epitope that is recognized and bound by the EP1 MoAb is obscured by ER conformational change and ER protein-protein interactions, while the target epitope for the H4624 MoAb remains accessible.

### Prediction of ER activity based on nuclear H4624/EP1 staining ratio

We proceeded to use the observed differences in staining behaviour between EP1 and H4624 MoAbs in cells with active ER to develop a dual staining assay to predict the activity state of ER in cells. Absolute nuclear staining intensity values are unlikely to be sufficiently predictive for ER pathway activity, due to for example variations in staining procedure or sample quality which may affect absolute fluorescent staining intensities. For that reason, per measured cell the ratio (H4624/EP1) was calculated between staining intensity obtained with the EP1 and the H4624 MoAbs. As can be seen the H4624/EP1 ratio increased in the ER active condition (Fig. 5D). Compared to individual MoAb staining (Fig. 5A,B), the H4624/EP1 ratio was better able to separate the ER-inactive condition from the ER-active condition (Fig. 5D). The expectation that the H4624/EP1 ratio would be superior in predicting ER activity, was confirmed by comparing its receiver operating characteristic (ROC) curve (AUC=0.87) with that of single MoAb EP1 (AUC=0.68) and H4624 (AUC=0.53) (Fig. 5E-G). The workflow steps leading to the H4624/EP1 ratio and ROC curve are illustrated in Supplementary Figs. 3 and 4. Similar results were obtained by replacing H4624 by the polyclonal antibody H184 (Supplementary Fig. 2C,D).

## Discussion

In two ER positive breast cancer cell lines, cell nuclear ER expression was measured by staining with various ER MoAbs, and compared with ER transcriptional activity measured with the mRNA-based ER pathway activity assay developed in our lab. This assay measures functional activity of the ER transcription factor as a readout for ER pathway activity^1^.

We show that nuclear presence of ER in a cell is not sufficient for ER transcriptional activity: cells stained positive for ER in the absence of a transcriptionally active ER. This confirms earlier results obtained in clinical studies, in which the ER pathway activity assay was performed on tissue samples from patients with ER positive breast cancer. In ER positive breast cancer, the ER pathway was shown to be either inactive or active, and an ER inactive pathway was associated with progressive disease under neoadjuvant endocrine therapy and with increased incidence of recurrence under adjuvant endocrine therapy^1^,^3^,^11^.

The estrogen receptor belongs to the nuclear receptor family of transcription factors and requires ligand-induced dimerization and recruitment of coactivator proteins to become transcriptionally active ^12^. Indeed, availability of estrogen ligand determined the activity status of nuclear ER, in line with the known mechanism of activation of the ER signaling pathway. While all used MoAbs stained nuclear ER, there were remarkable differences in staining behaviour: ER staining by EP1 and 1D5 MoAbs decreased rapidly upon transcriptional activation of ER, while staining with H4624 MoAb appeared to be independent of ER activity. We hypothesized that during the ER activation process, which is associated with conformational changes in ER proteins and binding to multiple other proteins to form an active transcription complex, the single binding epitope for the EP1 and 1D5 MoAbs becomes hidden and inaccessible for antibody binding, in contrast to the binding epitope for the H4624 MoAb (Fig. 5). Nuclear ER staining by the polyclonal ER antibody H184 was also independent of the activation state of ER, providing support for this hypothesis: only a limited number of the many binding epitopes for this mixture of antibodies is expected to become hidden upon ER activation.

The observation that nuclear ER staining using EP1 and 1D5 MoAbs differed depending on the activity state of ER, allowed development of a dual antibody staining assay for improved prediction of ER activity state. The described method requires generation of a digital image with a standard digital pathology scanner, application of a cell nucleus detection algorithm and *per cell* calculation of the nuclear H4624/EP1 fluorescence intensity ratio. The H4624/EP1 ratio score is correlated with, and predicts, the probability of ER activity. In principle the EP1 MoAb can be replaced by 1D5 or another MoAb of which the binding epitope becomes inaccessible upon ER activation. Similarly the H4624 MoAb can be replaced by the H184 polyclonal antibody or another antibody which binds ER independent of its activity state. Translation from a fluorescent to an immunohistochemistry-based assay should also be possible. The concept that underlies the dual staining assay, that is, use digitized staining data obtained with two antibodies which recognize different functional states of a protein to calculate a staining intensity ratio per cell, can in principle be applied to development of staining assays for many different proteins of which the functional state needs to be characterized.

When compared to the mRNA-based ER pathway activity assay, the dual staining ER activity assay has the advantage of being able to assess ER activity within preserved tissue architecture, but it lacks the quantitative nature of the mRNA-based assay. It needs to be explored to what extent the assay is compatible with other types of sample preparation, such as formalin fixated paraffin embedded tissue ^13^. The presented staining assay is expected to have potential value, complementary to the mRNA-based ER pathway activity assay. For example, when in a tissue sample the ER pathway is measured as active using the ER pathway assay, the dual ER staining assay provides information on type and location of the ER active cells in the tissue.

## Conclusion

Routine ER staining provides insufficient information on functional activity of ER. A dual ER staining assay was developed which improves ER activity assessment in cells on a scanned slide. This assay may find applications in preclinical research on the ER pathway, in development of ER activity-modifying drugs, as well as for clinical diagnostics. The concept that underlies this assay is expected to be generalizable to development of assays for other proteins of which characterization of the functional activity state on a cell or tissue slide is of value.

## Supplementary Methods

### Development of cell and nucleus recognition algorithms and nuclear ER staining quantification algorithm

Algorithms for analysis of digital images and quantification of immunofluorescent staining intensities were developed using MATLAB 2012b (The MathWorks Inc., Natick, MA, 2000, United States). Digital image analysis consisted of: (1) image pre-processing; (2) cell nucleus detection; (3) cell membrane location estimation; and (4) quantification of immunofluorescence staining intensity in detected nuclei. In the image pre-processing step the input image is subsampled by a factor of three to reduce the computational load for the subsequent steps. The image is smoothed lightly with a Gaussian smoothing filter with a sigma of 1 to suppress sensor noise. The nucleus detection step uses the DAPI signal detection channel and consists of three steps: dynamic thresholding, cleaning, and splitting. In the first step, for reasons of varying background intensity, the DAPI channel image is thresholded dynamically, meaning that the threshold is not a single value for the entire image but is adapted for background intensity. This is accomplished by strongly smoothing the DAPI image using a Gaussian smoothing filter with a sigma of 4 to flatten out nuclear fluorescent signals while preserving the much coarser variations in background intensity. The smoothed image serves as the (dynamic) threshold for the unsmoothed DAPI image resulting in a binary image that marks nuclei with a fluorescence intensity that exceeds the dynamic threshold. The second nucleus detection step, the cleaning step, consists of removing objects from the created binary image that clearly are not nuclei or clusters of nuclei, based on morphology criteria. The final nucleus detection step splits clustered nuclei into separate ones, based on identification of a local minimum along the nuclear fusion line. To find these lines, a distance transform was applied to the inverted binary nucleus image, followed by applying a watershed algorithm to the distance-transformed image. Nuclear fusion lines were used to split up clustered nuclei in separate nuclei. The membrane estimation step uses the detected nuclei to estimate the location of the cell membranes. Once nuclei had been identified, staining intensities were quantified for each imaged cell, using separate fluorescent readout channels for dual fluorescent antibody staining. For all experiments image scanning parameters (e.g. exposure time and gain) were kept constant, enabling quantitative comparison between staining intensities across different image scans.

### Supplement

**Supplementary Figure 1.**
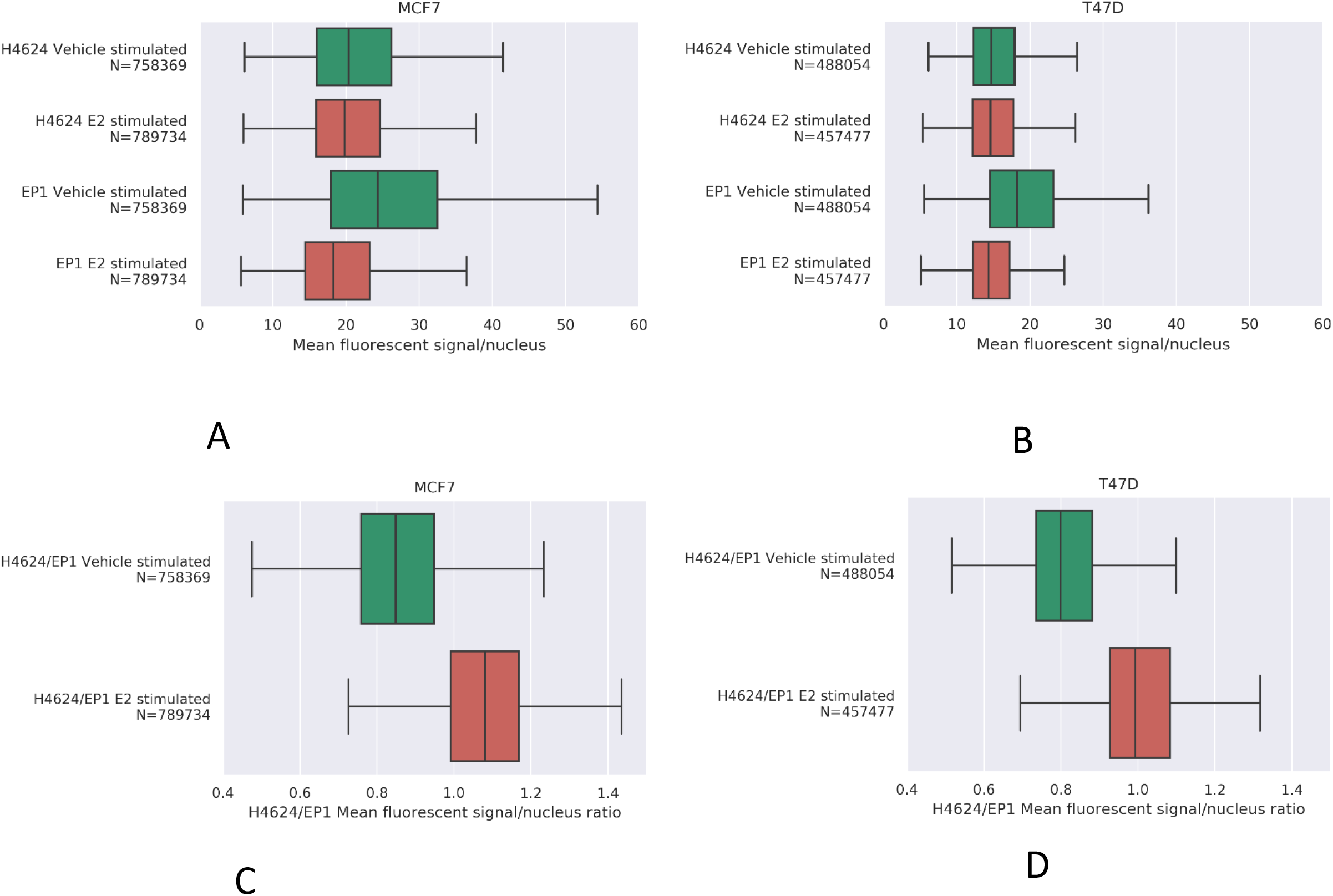
A,B. Comparison between nuclear immunofluorescent staining intensity with EP1 and H4624 MoAbs in MCF7 and T47D breast cancer cells, treated with vehicle or estradiol for 30 minutes. Boxplots of mean fluorescent signal intensity per cell nucleus for MCF7 cells (A) and T46D cells (B), stained with either EP1 or H4624 MoAb. C,D. Comparison of mean H4624/EP1 ratio between MCF7 (C) and T47D (D) cells. Depicted is the median, with box visualizing 2nd and 3rd quartile.

**Supplementary Figure 2:**
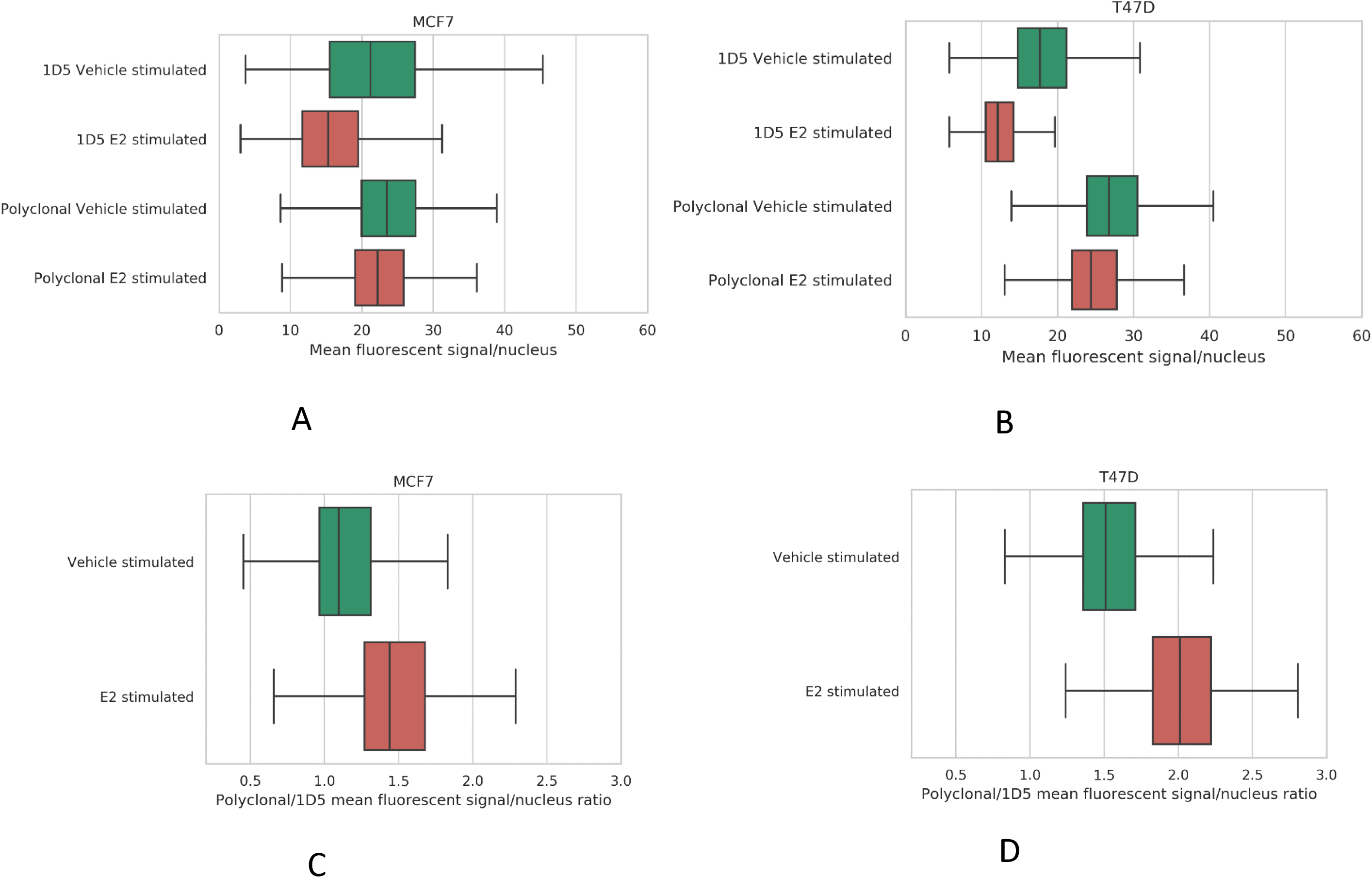
A,B. Comparison between nuclear immunofluorescent staining with 1D5 MoAb and polyclonal ER antibody in MCF7 and T47D breast cancer cells, estradiol-deprived and either treated with vehicle or with estradiol (E2) for 30 minutes. Boxplots of mean fluorescent signal intensity per cell nucleus for MCF7 cells (A) and T46D cells (B), stained with either 1D5 MoAb or polyclonal antibody H184. C,D. H184/1D5 ratio in ER inactive (vehicle) and ER active (E2 stimulated) MCF7 (C) and T47D (D) cells. Depicted is the median, with box visualizing 2nd and 3rd quartile.

**Supplementary Figure 3.**
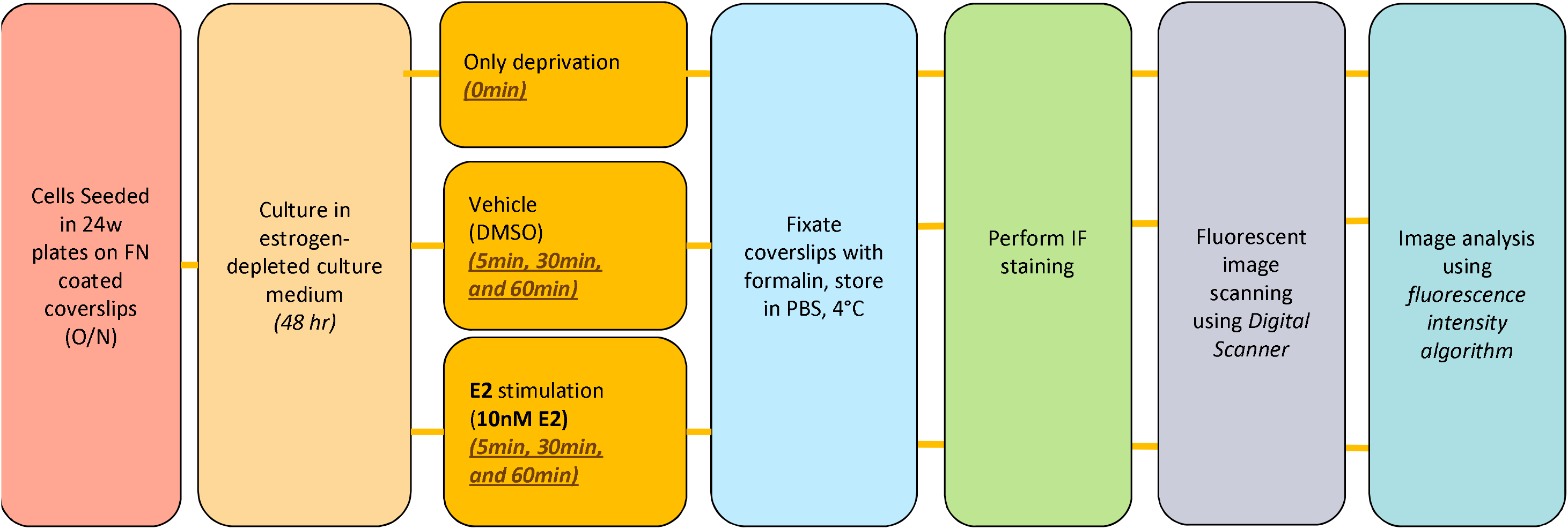
Estrogen deprivation/stimulation (E2) experiments with MCF7 and T47D cells on glass coverslips. Coverslips were coated with fibronectin (FN). IF: immunofluorescence; O/N: overnight.

**Supplementary Figure 4:**
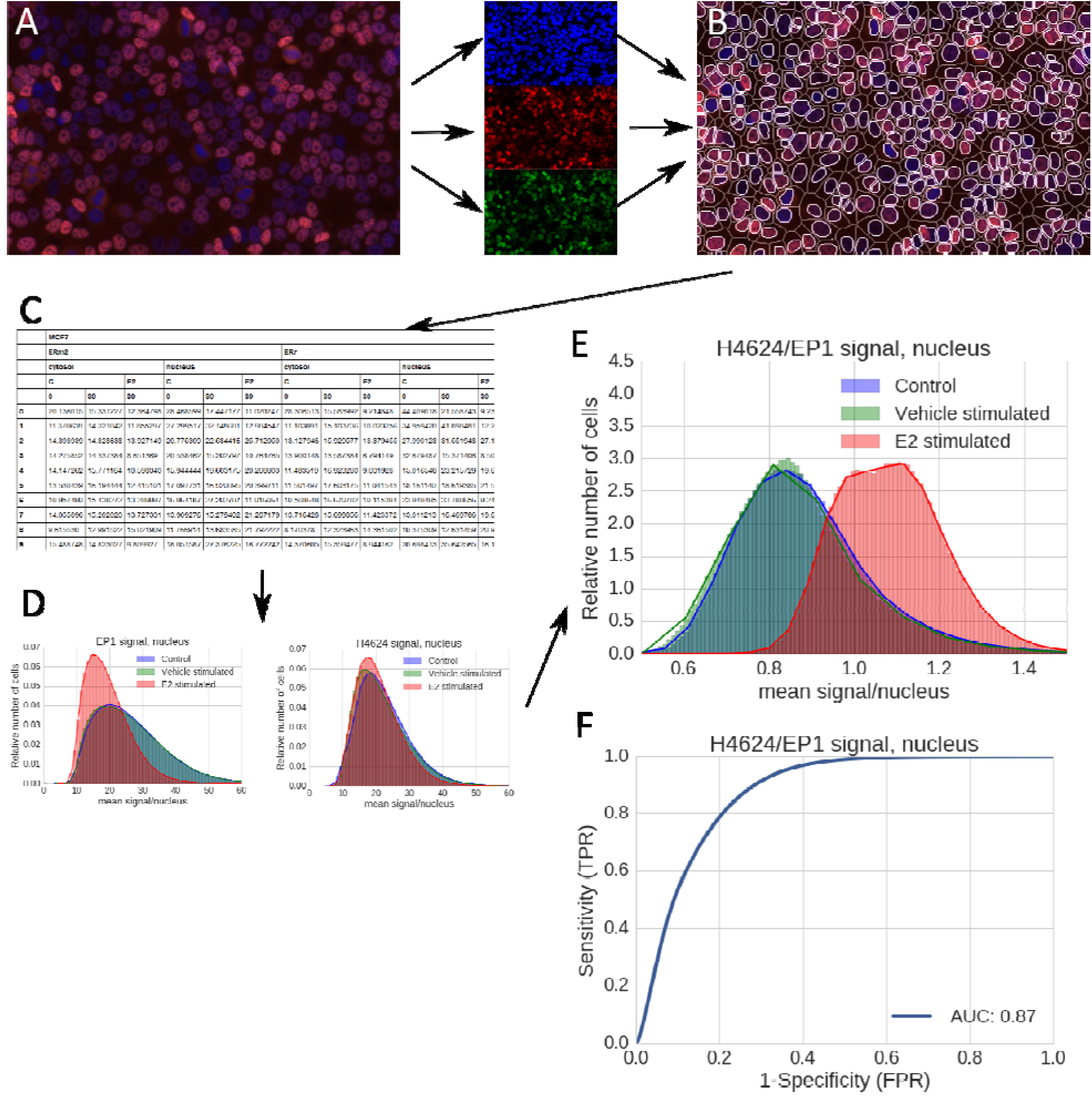
Dual immunofluorescent staining method: An image is taken with the Digital Scanner using 3 fluorescent channels, DAPI, FITC and Cy5 (A). The DAPI channel is used to identify all the cells and define a nuclear and cytosolic compartment (B) in which H4626-FITC and EP1-Cy5 signals are quantified (C). The EP1-Cy5 staining intensity reduces upon stimulation with E2 while the H4626-FITC staining intensity is unaffected (D). Nuclear H4626-FITC signal is used as a proxy for the total number of ER molecules present in the nucleus, irrespective of ER’s activation status; and the EP1-Cy5 signal is used as a proxy for the number of ER molecules that lose their EP1 binding epitope upon activation of the ER pathway. When dividing the H4626-FITC signal by the EP1-Cy5 signal, the obtained ratio is independent of the absolute amount of nuclear ER, and is hypothesized to represent the fraction of ER proteins which change conformation and available antibody-binding epitopes upon E2 binding and coactivator protein binding, and consequent ER transcriptional activation. The H4626-FITC/EP1-Cy5 ratio shows improved separation of cell populations with respectively an inactive and an active ER pathway (E). The ROC curve supports the ratio as a good indicator of ER activity (F).

**Supplemental Table 1.** Used primary and secondary antibodies.

| Used abbreviation | Antibody Name | Clone | Isotype (kDa) | Company/ Product # | Dilution (µg/mL) |
| --- | --- | --- | --- | --- | --- |
| Primary/Unlabeled antibodies: |  |  |  |  |  |
| ID5 | Monoclonal Mouse Anti-Human Estrogen Receptor $\alpha$ | 1D5<br>For clinical use | IgG1k | Dako/Cat#M7047<br>(0.2mL) | 1:60<br>(2.8µg/ml) |
| EP1 | Monoclonal rabbit anti-human estrogen receptor $\alpha$ | EP1<br>For clinical use | | Dako/Cat#M364329<br>(0.2mL) | 1:40 |
| H4624 | Estrogen receptor $\alpha$ mouse anti-human mAb | H4624<br>for research use only | IgG2a | Invitrogen /Cat#417000<br>(100µg) | 1:50 (20µg/ml) |
| H-184 | Polyclonal rabbit anti-human estrogen receptor- $\alpha$ | H-184 | | Polyclonal; 1/50 dilution, Santa Cruz Biotechnology, Dallas, Texas, USA | 1:50 |
| Secondary antibodies: |  |  |  |  |  |
| Goat anti-mouse | goat anti-mouse ALEXA-555 |  |  | Invitrogen cat nr A21422 | 1:200 |
| Goat anti-rabbit | goat anti-rabbit IgG ATTO-633 |  |  | Sigma cat nr 41176-1ML | 1:200 |

